# Adenovirus-Vectored SARS-CoV-2 Vaccine Expressing S1-N Fusion Protein

**DOI:** 10.1101/2022.05.09.491179

**Authors:** Muhammad S. Khan, Eun Kim, Alex McPherson, Florian J. Weisel, Shaohua Huang, Thomas W. Kenniston, Elena Percivalle, Irene Cassaniti, Fausto Baldanti, Marlies Meisel, Andrea Gambotto

## Abstract

Additional COVID-19 vaccines that are safe, easy to manufacture, and immunogenic are needed for global vaccine equity. Here, we developed a recombinant type 5 adenovirus vector encoding for the SARS-CoV-2-S1 subunit antigen and nucleocapsid as a fusion protein (Ad5.SARS-CoV-2-S1N) delivered to BALB/c mice through multiple vaccine administration routes. A single subcutaneous (S.C.) immunization with Ad5.SARS-CoV-2-S1N induced a similar humoral response, along with a significantly higher S1-specific cellular response, as a recombinant type 5 adenovirus vector encoding for S1 alone (Ad5.SARS-CoV-2-S1). Immunogenicity was improved by homologous prime boost strategies, using either S.C. or intranasal (I.N.) delivery of Ad5.SARS-CoV-2-S1N, and further improved through heterologous prime boost, with traditional intramuscular (I.M.) injection, using subunit recombinant S1 protein. Priming with low dose (1×10^10^ v.p.) of Ad5.SARS-CoV-2-S1N and boosting with either wildtype recombinant rS1 or B.1.351 recombinant rS1 induced a robust neutralizing response, that was sustained against immune evasive Beta and Gamma SARS-CoV-2 variants, along with a long-lived plasma cell response in the bone marrow 29 weeks post vaccination. This novel Ad5-vectored SARS-CoV-2 vaccine candidate showed promising immunogenicity in mice and supports the further development of COVID-19 based vaccines incorporating the nucleoprotein as a target antigen.

## Introduction

The ongoing COVID-19 pandemic, caused by Severe Acute Respiratory Syndrome Coronavirus 2 (SARS-CoV-2), continues to have a large impact on public health across the globe [1–3]. COVID-19 first emerged in 2019, was declared a global pandemic by the World Health Organization on March 11, 2020 and has since claimed approximately 5.6 million lives as of January 24, 2022. Vaccination continues to be one of the most effective public health interventions to curb infectious diseases and their impact on human, and animal, health [4–7]. Currently approved COVID-19 vaccines have been a key tool in fighting this pandemic, however, they have been hampered by worldwide distribution inequalities that have left many low to middle income countries without access [8–13]. With many countries now distributing a COVID-19 booster to those already vaccinated, global vaccine inequality is at risk of increasing further [14–17]. Global vaccine inequality has, and will continue to, lead to new SARS-CoV-2 variants that may be able to escape natural and vaccine acquired immunity [18–21]. To this effect, new COVID-19 vaccines are needed, that are better suited for worldwide vaccination and that incorporate strategies to target more conserved regions of SARS-CoV-2, to combat new viral variants.

Beta coronaviruses (Beta-CoVs), such as SARS-CoV-2, are enveloped, positive sense, ssRNA viruses [22, 23]. BetaCoVs encode the envelope, nucleocapsid (N), membrane, and spike (S) proteins [24, 25]. The spike protein has been the focus of currently approved COVID-19 vaccines, and of various COVID-19 vaccines in development, due to its role in viral infection of host cells [26]. The S protein comprised of two subunits, S1 and S2, that allow for viral attachment to the host cell receptor and in fusion to the host cellular membrane [26, 27]. It has been demonstrated that antibodies targeting the S protein can block the binding of SARS-CoV-2 to the cell receptor, allowing the S protein to be a focal target of vaccine development [28–32]. Our previously published reports on vaccines against not only SARS-CoV-2, but also SARS-CoV-1 and MERS, have shown the ability of S1 subunit targeting vaccines to generate neutralizing antibody response [33–36]. Our previous efforts have established the immunogenicity of S1 based Beta-CoVs vaccines through both Adenoviral (Ad) -vectored vaccines and subunit recombinant protein vaccines. Most recently we have described an Ad-vectored SARS-CoV-2 vaccine expressing S1 alone generating a robust immune response in BALB/cJ mice [34]. We have also presented skin-targeted S1 subunit protein vaccines that induce antigen specific antibody responses against MERS-CoV and SARS-CoV-2 [33].

Of particular interest is the investigation of including more conserved regions of SARS-CoV-2, such as the N protein, in vaccine strategies to combat emerging variants. The primary function of the SARS-CoV-2 N protein is to package the viral genome [37].The N protein is more conserved than the S protein, with 90% amino acid homology, and also accumulates fewer mutations over time [38, 39]. The N protein also contains key T cell epitopes for SARS-CoV-2 immunity [40–42]. Taken together, the literature suggests potential for an S1; and N; based COVID-19 vaccine that incorporates the neutralizing antibody response to S1 with the conserved T cell response to N.

Adenovirus (Ad)-vectored vaccines have been investigated thoroughly due to their ability to induce a balanced humoral and cellular immune response [43–45]. Our previous studies using Ad vectored vaccines expressing SARS-CoV-2-S1, SARS-CoV-1-S1, and MERS-S1 have illustrated the potential for Ad-based vaccines against coronavirus diseases. Indeed, there have been a number of Ad based SARS-CoV-2 (including Ad5, Ad26, and chimpanzee adenovirus vectors) vaccines that have shown promising results in clinical trials [46]. The Ad5-vectored COVID-19 vaccine, CanSino Convidicea Vaccine (Ad5-nCoV), encoding for the full S protein has also shown promising immunogenicity when delivered intramuscularly and has been approved by multiple countries [47–49]. Although there has been remarkable progress of Ad-based COVID- 19 vaccines, there is still a need for investigation of novel vaccine strategies, such as including the N protein, or homologous and heterologous prime-boost strategies, that induce sustained immunity against SARS-CoV-2 variants.

Here, we describe the development of multiple Ad-vectored and subunit recombinant protein SARS-CoV-2 vaccine candidates against COVID-19. We designed and constructed a recombinant type 5 Ad vector encoding for a fusion protein S1N subunit antigen (Ad5.SARS-CoV-2-S1N). We evaluated the immunogenicity of a single shot, along with homologous and heterologous prime boost, of this vaccine in BALB/cJ mice through multiple delivery routes including intranasal delivery (I.N.), subcutaneous injection (S.C.), and intramuscular injection (I.M.). We investigated the ability of Ad5.SARS-CoV-2-S1N to induce antigen-specific humoral and cellular immune responses and investigated the virus-specific neutralization activity of the generated antibodies. Our study demonstrates the development of an Ad-based SARS-CoV-2 vaccine, along with heterologous boost using subunit recombinant SARS-CoV-2 vaccine, that can induce robust and durable SARS-CoV-2 specific immune response in mice, which is sustained against Beta (B.1.351) and Gamma (P.1) SARS-CoV-2 variants.

## Results

### Adenoviral SARS-CoV-2-S1N Vaccine

To produce E1/E3 deleted replication-deficient human type 5 adenovirus expressing SARS-CoV-2-S1 protein, we generated pAd/SARS-CoV-2-S1N by subcloning the codon-optimized SARS-CoV-2-S1 & wild-type Nucleoprotein gene into the shuttle vector, pAd (GenBank U62024) at SalI & NotI sites. Next, Ad5.SARS-CoV-2-S1N (Ad5.S1N) was created by loxP homologous recombination (**Fig. 1A**). To detect SARS-CoV-2-S1 expression driven by the generated Adenoviral candidate, the serum-free supernatants from A549 cells infected with Ad5.S1N were characterized by SDS- PAGE and Western blot analysis. The recombinant SARS-CoV-2-S1N proteins (Ad5.S1N infected cells) were recognized by the polyclonal S1 and N antibody at the expected glycosylated monomer molecular weights of about 150 kDa (**Fig. 1B**, **Fig. 1C**).

**Figure 1.**
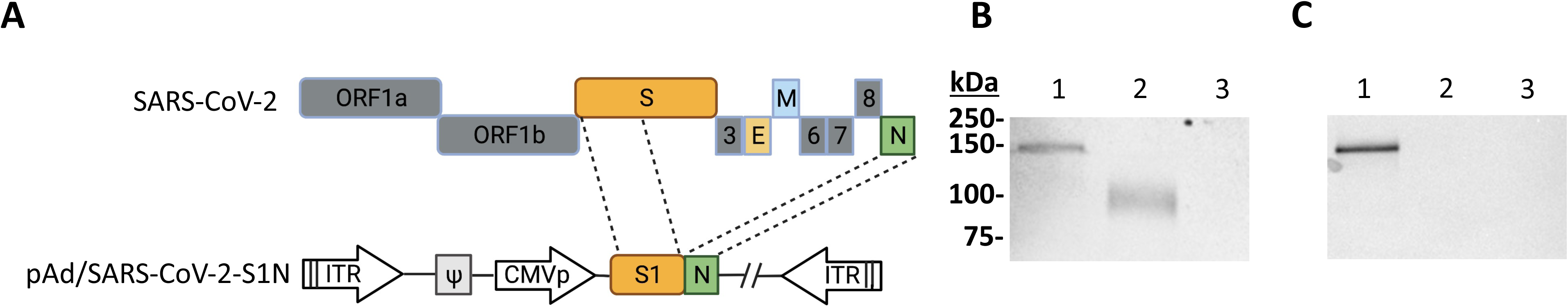
Adenoviral vectored SARS-CoV-2-S1N vaccine. **(A)** A shuttle vector carrying the codon optimized SARS-CoV-2-S1 gene encoding N terminal 1–661 along with full Nucleoprotein was designed as shown in the diagram. The vector was used to generate recombinant type 5 replication deficient adenoviruses (Ad5) by homologous recombination with the adenoviral genomic DNA. ITR, inverted terminal repeat. **(B)** Detection of the SARS-CoV-2-S1N fusion protein by western blot with the supernatant of A549 cells infected with Ad5.SARS-CoV-2.S1N (Ad5.S1N) using S1 SARS CoV-2 rabbit polyclonal antibody (lane 1). As a positive control, supernatant of A549 cells infected with Ad5.SARS-CoV-2.S1 (Ad5.S1) was loaded (lane 2). As a negative control, supernatant of A549 cells infected with an empty vector (AdΨ5) was loaded (lane 3). **(C)** Detection of the SARS-CoV-2-S1N fusion protein by western blot with the supernatant of A549 cells infected with Ad5.SARS-CoV-2.S1N (Ad5.S1N) using N SARS CoV-2 rabbit polyclonal antibody (lane 1). As a negative control, supernatant of A549 cells infected with Ad5.SARS-CoV-2.S1 (Ad5.S1) was loaded (lane 2). As a negative control, supernatant of A549 cells infected with an empty vector (AdΨ5) was also loaded (lane 3). The supernatants were resolved on SDS 10% polyacrylamide gel after being boiled in 2% SDS sample buffer with β-ME.

### Single shot immunogenicity of Adenoviral SARS-CoV-2-S1N vaccine

To evaluate the impact of including the N protein of SARS-CoV-2 in vaccine formulation through a S1N fusion protein, we first compared the immunogenicity of Ad5.SARS-CoV-2-S1N (Ad5.S1N) with our previously published Ad5.SARS-CoV-2-S1 (Ad5.S1), which expresses S1 alone [34]. We determined S1 specific IgG, IgG1, and IgG2a endpoint titers in the sera of vaccinated BALB/cJ mice (either via I.N. delivery or S.C. injection) and control mice (AdΨ5 immunized groups). Serum samples, collected from mice at the timepoints indicated, were serially diluted to determine SARS-CoV-2-S1 specific IgG, IgG1, and IgG2a endpoint titers for each immunization group using ELISA (**Fig. 2**). Results suggest that both Ad5.S1 and Ad5.S1N resulted in comparable S1-specific IgG, IgG1, and IgG2a endpoint titers (**Fig. 2**). S.C. injection of Ad5.S1 and Ad5.S1N resulted in significantly increased S1-specific IgG endpoint titers as early as 2 weeks after a single vaccination, compared to the mock vaccinated group (**Fig. 2A**, p < 0.05, Kruskal-Wallis test, followed by Dunn’s multiple comparisons). I.N. Ad5.S1 and Ad5.S1N SARS-CoV-2-S1 specific IgG endpoint titer increased to comparable levels of S.C. injection by week 3 and were sustained through week 8 (**Fig. 2A**). Both Ad5.S1 and Ad5.S1N, regardless of immunization delivery route, resulted in similar IgG1 and IgG2a endpoint titers indicating a balanced Th1/Th2 response (**Fig. 2B**). To evaluate the functional quality of vaccine-generate antigen-specific antibodies, we used a microneutralization assay (NT_90_) to test the ability of sera from immunized mice to neutralize the infectivity of SARS-CoV-2. Sera, collected from all mice 6 and 8 weeks after vaccination, were tested for the presence of SARS-CoV-2-specific neutralizing antibodies (**Fig. 2C**). As expected, there were no detected neutralizing antibody response in sera of AdΨ5 immunized mice. Neutralizing antibodies were detected in Ad5.S1 and Ad5.S1N vaccinated mice (for both I.N. and S.C. delivery routes) at week 6. Neutralizing antibody response for I.N. delivery was greater in magnitude for both Ad5.S1 and Ad5.S1N, when compared to S.C. injection, with S.C. injected mice having low to undetectable neutralizing antibodies by week 8 (**Fig. 2C**). Next, we evaluated the antigen-specific cellular immune response induced by the single immunization. We investigated S1 and N-specific cellular immunity in mice after vaccination by quantifying IFN-γ^+^ CD8^+^ and CD4^+^ T-cell responses through intracellular cytokine staining (ICS) and flow cytometry. Results suggest that I.N. delivery of either Ad5.S1 or Ad5.S1N did not induce increased systemic S1-specific or N-specific CD8^+^ T cell immunity, when compared to I.N. AdΨ5 control groups (**Fig. 3A**). However, S.C. vaccination induced increased systemic S1-specific IFN-γ^+^ CD8^+^ T cell responses for both Ad5.S1 and Ad5.S1N. S.C. vaccination of Ad5.S1 and Ad5.S1N resulted in significantly increased S1-specific IFN-γ^+^ CD8^+^ T cell response when compared to both I.N. AdΨ5 and S.C.

**Figure 2.**
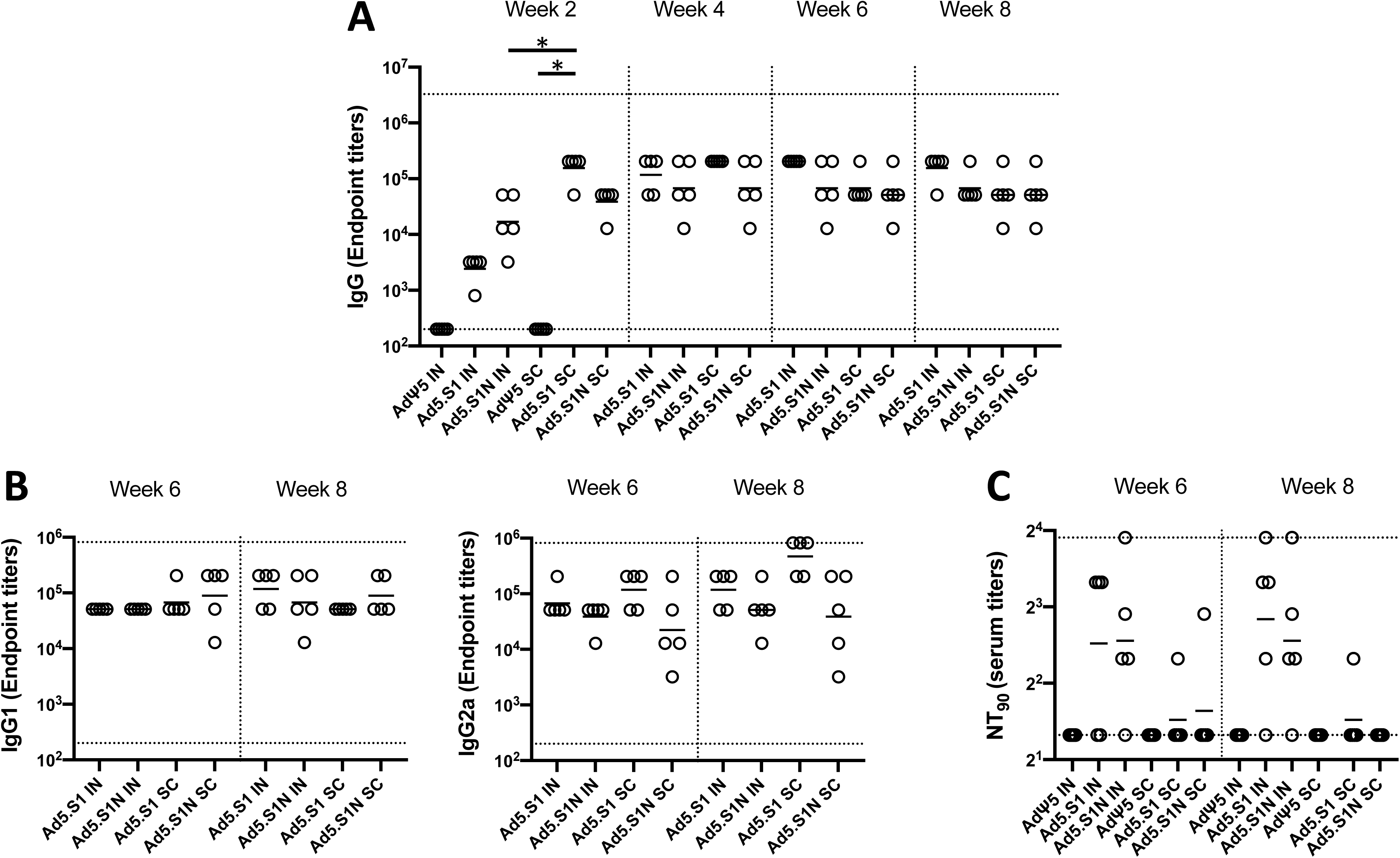
Antigen-specific antibody responses in mice immunized with adenoviral vectored SARS-CoV-2-S1N vaccine. BALB/c mice (n =5 mice per groups) were immunized intranasally (IN) or subcutaneously (SC) with 5×10^10^ v.p. of Ad5.SARS-CoV-2-S1N (Ad5.S1N), Ad5.SARS- CoV-2-S1 (Ad5.S1), or empty Ad5 vector as negative control (AdΨ5). On weeks 2, 4, 6 and 8 after vaccination, the sera from mice were collected, serially diluted (200x), and tested for the presence of SARS-CoV-2-S1-specific **(A)** IgG, **(B)** IgG1 & IgG2 antibody levels at the indicated time points by ELISA. **(C)** Serum from immunized mice was tested neutralizing antibodies using a plaque reduction neutralization test (PRNT) SARS-CoV-2 strain from Wuhan. Serum titers that resulted in a 90% reduction in SARS-CoV-2 viral plaques (NT90) compared to the virus control are reported at six and eight weeks after immunization, respectively, and bars represent geometric means. No neutralizing antibodies were detected in serum AdΨ5 control group. Significance was determined by Kruskal-Wallis test followed by Dunn’s multiple comparisons (\**p* < 0.05). Horizontal solid lines represent geometric mean antibody titers. Horizontal dotted lines represent minimum and maximum dilutions. Results are from a single animal experiment. (N=5 mice per group). ELISA experiments were conducted twice while neutralizing antibody experiments were conducted once.

**Figure 3.**
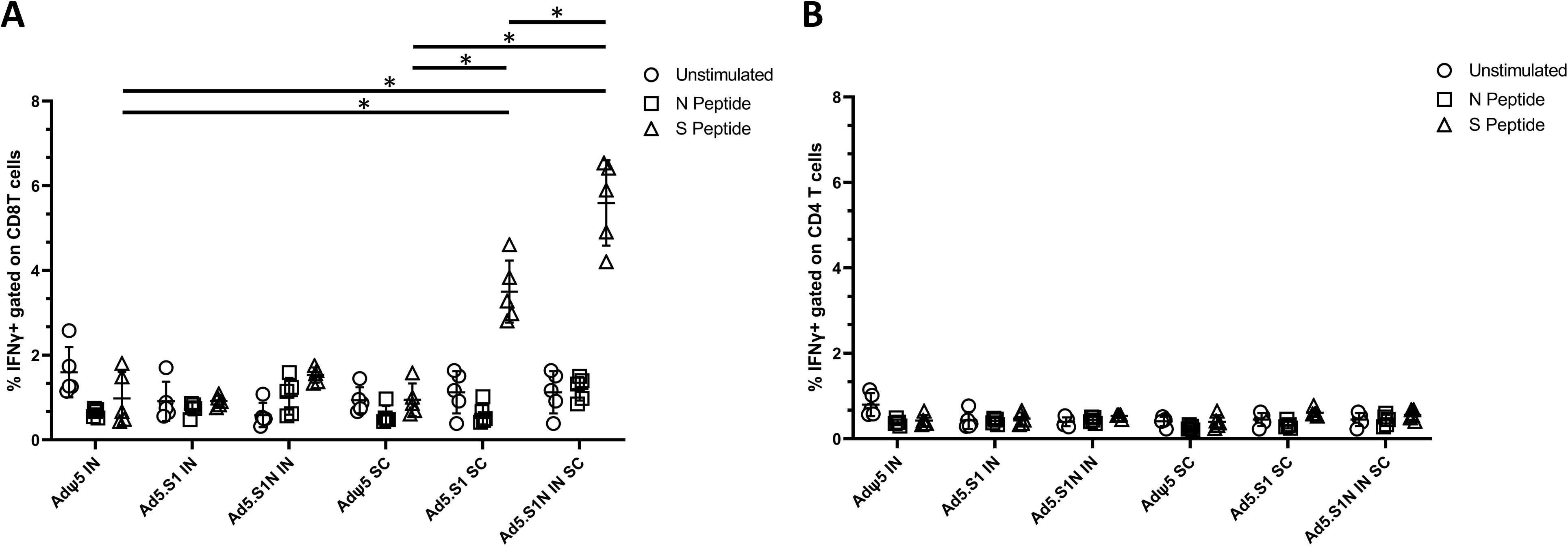
Antigen specific cellular responses in BALB/c mice immunized with Ad5.SARS-CoV-2-S1 and Ad5.SARS-CoV-2-S1N. BALB/c mice were immunized S.C. or I.N. with 5×10^10^ v.p. of Ad5.SARS-CoV-2-S1 (Ad5.S1), Ad5.SARS-CoV-2-S1N (Ad5.S1N) or AdΨ5. Ten days after vaccination, splenocytes were isolated and stimulated with SARS-CoV-2-S1 and SARS-CoV-2 Nucleoprotein PepTivator, followed by intracellular cytokine staining (ICS) and flow cytometry to identify SARS-CoV-2-S1 & Nucleoprotein specific T cells (see Supporting information for complete gating strategy). **(A)** Frequencies of SARS-CoV-2-S1 and Nucleoprotein specific CD8+ IFN γ^+^. Results are mean ± SD. **(B)** Frequencies of SARS-CoV-2-S1 and Nucleoprotein specific CD4^+^ IFN γ^+^. Groups were compared by one-way Welch’s ANOVA, followed by Dunnett’s T3 multiple comparisons, and significant differences are indicated by *p < 0.05. Unstimulated controls are represented by circles, Nucleoprotein stimulated samples (N peptide) are represented by squares, and S1 peptide stimulated samples (S peptide) are represented by triangles. Results are from a single animal experiment. (N = 5 mice per group). This experiment was conducted once.

AdΨ5 (**Fig. 3A**, p < 0.05, one-way Welch’s ANOVA, followed by Dunnett’s T3 multiple comparisons). S1-specific IFN-γ^+^ CD8^+^ T cell response for S.C. vaccinated Ad5.S1N mice were significantly higher than in S.C. vaccinated Ad5.S1 mice (**Fig. 3A**, p < 0.05, one-way Welch’s ANOVA, followed by Dunnett’s T3 multiple comparisons). However, S.C. vaccinated Ad5.S1N mice did not have significantly higher N-specific IFN-γ^+^ CD8^+^ T cell response when compared to S.C. vaccinated Ad5.S1N (**Fig. 3A**, p > 0.05, one-way Welch’s ANOVA, followed by Dunnett’s T3 multiple comparisons). There was no significant increase in S1-specific or N-specific IFN-γ^+^ CD4^+^ T cell response in vaccinated groups when compared to controls (**Fig. 3B**, p > 0.05, one-way Welch’s ANOVA, followed by Dunnett’s T3 multiple comparisons). In sum, these finding indicate that a single vaccination with Ad5.S1 or Ad5.S1N, via either S.C. injection or I.N. delivery, results in a robust S1-specific IgG response, with a balanced IgG1/IgG2a ratio. The choice of the route of vaccine administration (S.C. or I.N.) influences both SARS-CoV-2 neutralizing antibodies and S1-specific CD8^+^ T-cell responses. Most importantly, inclusion of the N protein, through Ad5.S1N, results in a significantly higher induction of S1-specific CD8^+^ T-cells when compared to Ad5.S1.

### Immunogenicity of homologous and heterologous prime boost vaccination strategy

To evaluate the ability to further increase neutralizing antibody and cellular immune response to vaccination with Ad5.S1N, we investigated prime boost strategies leveraging homologous and heterologous strategies. Prime boost vaccination strategies have been shown to increase the quantity, durability, and quality of the immune response to vaccination and have been employed for numerous types of vaccines [50, 51]. **Figure 4A** outlines the prime boost vaccination strategies used. Homologous prime and boost of Ad5.S1N using differing vaccination routes (I.N. and S.C.) was done to circumvent Ad5 vector immunity induced by prime immunization, which has been shown to limit the effectiveness of Ad5 vectored vaccines [52, 53]. We also evaluated a heterologous prime boost group that were primed S.C. with Ad5.S1N and boosted with 15ug of recombinant subunit S1 WT protein. Heterologous prime boost strategies have been shown to be more immunogenic than homologous prime boost regimens, necessitating its investigation [51, 54–56].

**Figure 4.**
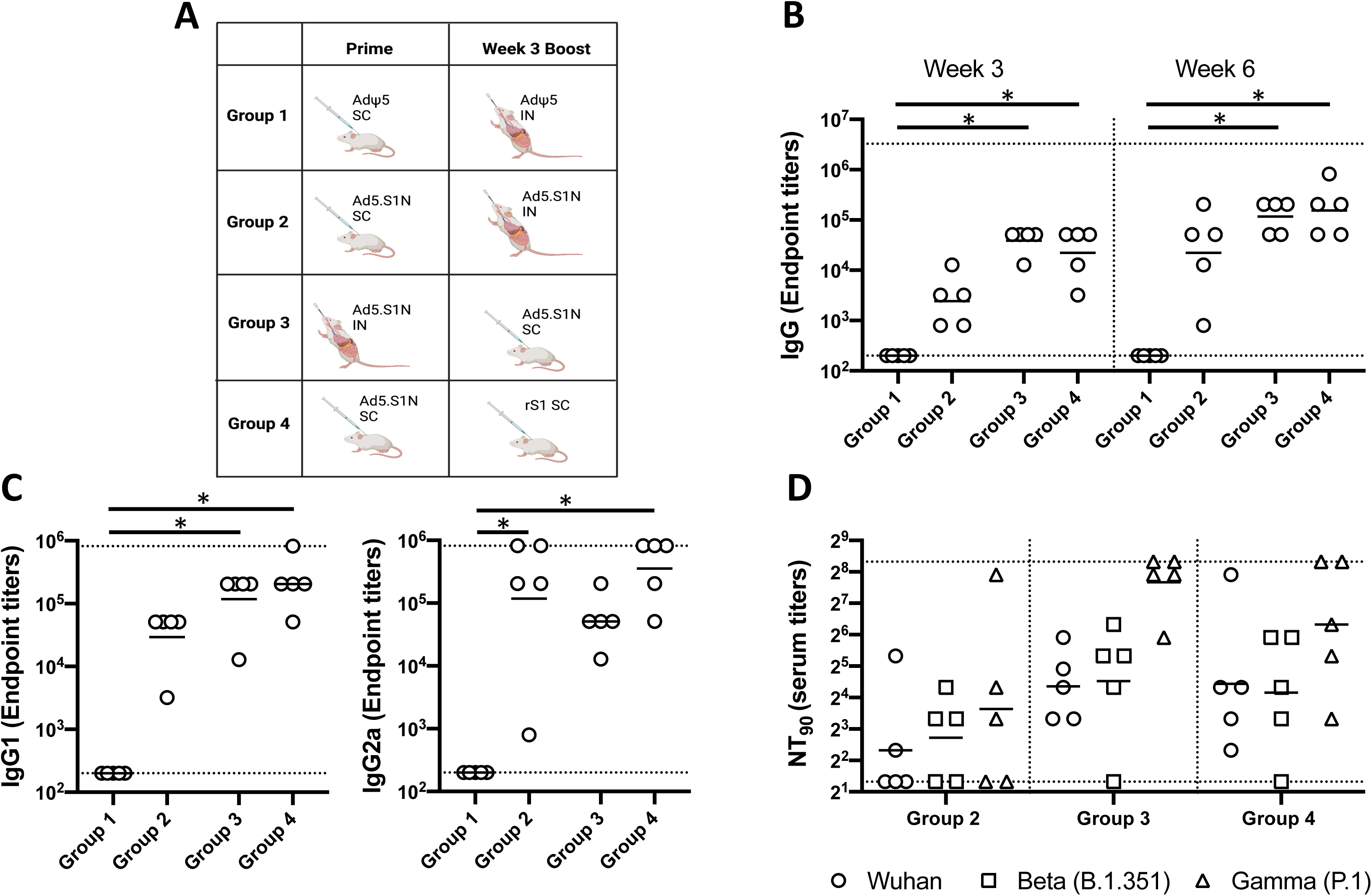
Homologous and heterologous prime boost vaccination with Ad5.SARS-CoV-2- S1N (Ad5.SIN) using differing vaccination routes along with recombinant S1 protein (rS1). **(A)** Schematic layout of mice (N = 5 mice per group) vaccinations. Group 1 served as control with SC AdΨ5 (1×10^10^ v.p.) prime and IN AdΨ5 (1×10^10^ v.p.) boost. Group 2 primed with SC Ad.S1N (1×10^10^ v.p.) and IN Ad.S1N (1×10^10^ v.p.) boost. Group 3 primed with IN Ad.S1N (1×10^10^ v.p.) and SC Ad.S1N boost (1×10^10^ v.p.). Group 4 primed with SC Ad.S1N (1×10^10^ v.p.) and SC rS1 boost (15ug). On weeks 3 and 6 after vaccination, the sera from mice were collected, serially diluted (200x), and tested for the presence of SARS-CoV-2-S1-specific **(B)** IgG antibody levels. **(C)** Week 6 sera was tested for presence of IgG1 and IgG2 antibody levels. **(D)** Week 6 serum from immunized mice was tested for neutralizing antibodies using a plaque reduction neutralization test (PRNT) with three different SARS-CoV-2 strains from Wuhan, South Africa (Beta B.1.351), or Brazil (Gamma P.1). Neutralization of Wuhan strain represented by circle, neutralization of Beta B.1.351 represented by square, and neutralization of Gamma P.1 represented by triangle. Serum titers that resulted in a 90% reduction in SARS-CoV-2 viral plaques (NT90) compared to the virus control are reported at six weeks after immunization and bars represent geometric means. No neutralizing antibodies were detected in serum PBS control group (not shown). Significance was determined by Kruskal-Wallis test followed by Dunn’s multiple comparisons (\**p* < 0.05). Horizontal lines represent geometric mean antibody titers. Horizontal dotted lines represent minimum and maximum dilutions. Results are from a single animal experiment. (N=5 mice per group). ELISA experiments were conducted twice while neutralizing antibody experiments were conducted once.

We determined S1 specific IgG, IgG1, and IgG2a endpoint titers in the sera of prime boost vaccinated BALB/cJ mice (either via I.N. delivery or S.C. injection) and control mice (AdΨ5 immunized groups). Serum samples, collected from mice at the timepoints indicated, were serially diluted to determine SARS-CoV-2-S1 specific IgG, IgG1, and IgG2a endpoint titers for each immunization group using ELISA (**Fig. 4**). Interestingly, I.N. prime and S.C. boost Ad5.S1N (Group 3), along with Ad5.S1N S.C prime and S.C. rS1 boost (Group 4), mice resulted in significantly higher IgG at weeks 3 and 6 when compared to control group (Group 1), while S.C. prime and S.C. boost Ad5.S1N (Group 2) did not (**Fig. 4B**, p < 0.05, Kruskal-Wallis test, followed by Dunn’s multiple comparisons). IgG1 and IgG2a endpoint titers were relatively balanced for experimental groups, however, Group 3 and Group 4 IgG1 endpoint titers were statistically significant when compared to Group 1; while Group 2 and Group 4 IgG2a endpoint titers were statistically significant when compared to Group 1 (**Fig. 4C**, p < 0.05, Kruskal-Wallis test, followed by Dunn’s multiple comparisons). We used a microneutralization assay (NT_90_) to test the ability of sera from immunized mice to neutralize the infectivity of SARS-CoV-2. The ability of sera to neutralize Beta (B.1.351) and Gamma (P.1) variants was also investigated. Sera, collected from all mice at 6 weeks after prime vaccination, were tested for the presence of SARS-CoV-2-specific neutralizing antibodies (**Fig. 4D**). Neutralizing antibodies were detected in in Group 2, Group 3, and Group 4, that was sustained, or increased, against Beta and Gamma variants. Group 2 had the lowest neutralizing antibody response when compared to Group 3 and Group 4 (**Fig. 4D**). Prime boost vaccination resulted in approximately 3-fold greater NT_90_ serum titers when compared to single shot immunization. We then evaluated the antigen-specific cellular immune response induced by homologous and heterologous prime boost immunization. We investigated S1-specific and N-specific cellular immunity in Group 1, Group 2, and Group by quantifying IFN-γ^+^ CD8^+^ and CD4^+^ T-cell responses through intracellular cytokine staining (ICS) and flow cytometry 8 weeks post prime vaccination. Results suggest that route of both prime and boost vaccination has an impact on both systemic S1-specific and N-specific CD8^+^ T cell immunity. Although, S.C. and I.N. boost of Ad5.S1N resulted in S1-specific CD8^+^ T cell immunity, this was not significant when compared to empty vector control (**Fig. 5A**, p > 0.05, one-way Welch’s ANOVA, followed by Dunnett’s T3 multiple comparisons). No N-specific CD8^+^ T cell immunity was detected for S.C. prime and I.N. boost of Ad5.S1N vaccinated mice. However, I.N. prime and S.C. boost of Ad5.S1N resulted in systemic S1-specific CD8^+^ T cell immunity with S1-specific CD8^+^ IFN-γ^+^ amounts being significantly higher when compared to empty vector control (**Fig. 5A**, p < 0.05, one-way Welch’s ANOVA, followed by Dunnett’s T3 multiple comparisons). While there was an induction in both S1-specific and N-specific IFN-γ^+^ CD4^+^ T cell response in I.N. prime and S.C. boost of Ad5.S1N vaccinated groups, it was not statistically significant when compared to controls (**Fig. 5B**, p > 0.05, one-way Welch’s ANOVA, followed by Dunnett’s T3 multiple comparisons). Taken together, these findings indicate that prime boost vaccination with Ad5.S1N results in higher S1-specific IgG, IgG1, IgG2a, and neutralizing response when compared to single immunization alone. The neutralizing ability was robust with I.N. prime and S.C. boost of Ad5.S1N, along with S.C. Ad5.S1N prime and S.C. rS1 boost. Prime boost vaccination of Ad5.S1N indicated the choice of vaccine administration order (S.C. or I.N.) impacted S1-specific and N-specific CD8^+^ T cell response.

**Figure 5.**
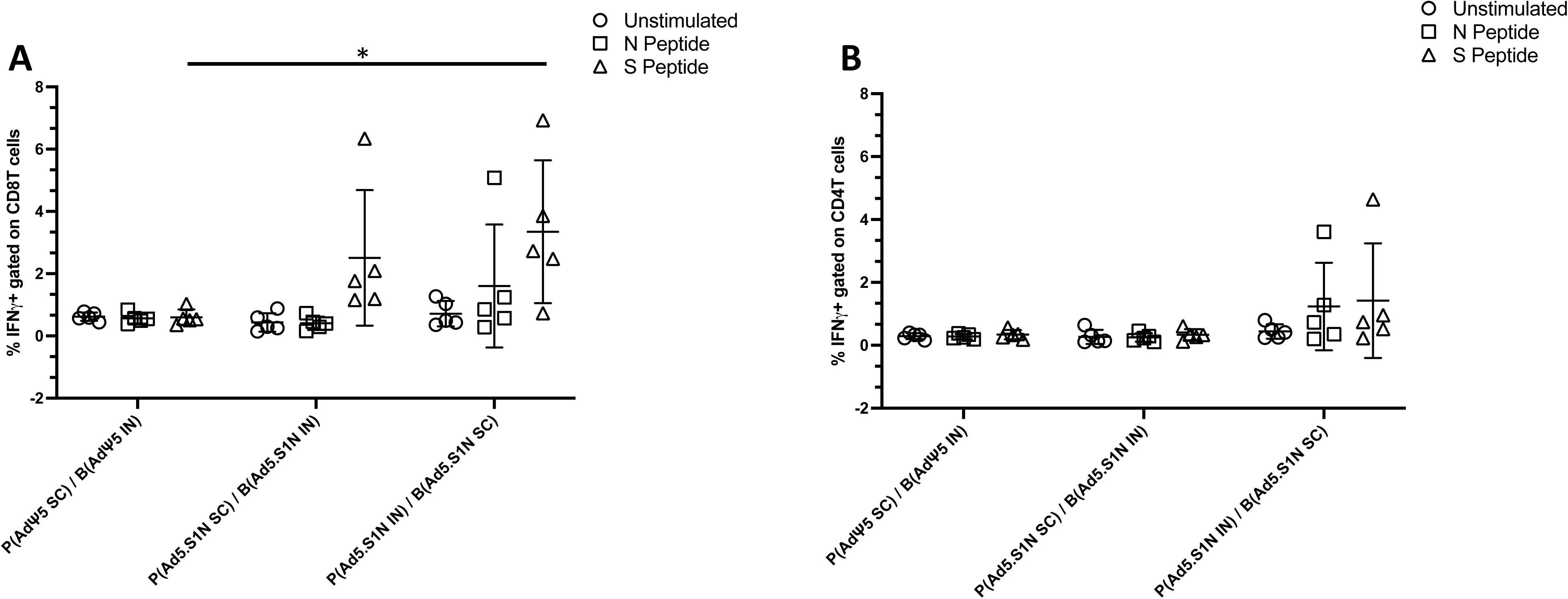
Antigen-specific cellular responses in BALB/c mice heterologous prime-boost immunized with Ad5.SARS-CoV-2-S1N. BALB/c mice were immunized as follows: Group 1 served as control with SC AdΨ5 (1×10^10^ v.p.) prime and IN AdΨ5 (1×10^10^ v.p.) boost. Group 2 prime SC Ad.S1N (1×10^10^ v.p.) and IN Ad.S1N (1×10^10^ v.p.) boost. Group 3 prime IN Ad.S1N (1×10^10^ v.p.) and SC Ad.S1N (1×10^10^ v.p.) boost. 8 weeks after initial vaccination, 5 weeks post boost, splenocytes were isolated and stimulated with SARS-CoV-2-S1 and SARS-CoV-2 Nucleoprotein PepTivator, followed by intracellular cytokine staining (ICS) and flow cytometry to identify SARS-CoV-2-S1 & Nucleoprotein specific T cells (see Supporting information for complete gating strategy). **(A)** Frequencies of SARS-CoV-2-S1 and Nucleoprotein specific CD8+ IFN γ^+^. **(B)** Frequencies of SARS-CoV-2-S1 and Nucleoprotein specific CD4^+^ IFN γ^+^. Results are mean ± SD. Groups were compared by one-way Welch’s ANOVA, followed by Dunnett’s T3 multiple comparisons, and significant differences are indicated by *p < 0.05. Unstimulated controls are represented by circles, Nucleoprotein stimulated samples (N peptide) are represented by squares, and S1 peptide stimulated samples (S peptide) are represented by triangles. Results are from a single animal experiment. (N = 5 mice per group). This experiment was conducted once.

### Immunogenicity of heterologous prime boost vaccination with Ad.S1N and variant specific recombinant S1 subunit protein

Due to the robust antibody response to S.C. Ad5.S1N prime and S.C. rS1 boost in Figure 5, we aimed to further compare the immunogenicity of recombinant monomeric S1 subunit vaccine formulation, at a higher dose of 45ug, with Ad5.S1N. **Figure 6A** outlines the prime boost vaccination strategies investigated with prime boost of 45ug rS1 WT, rS1 B.1.351, a combination of rS1 WT & rS1 B.1.351, and Ad5.S1N. A key advantage of harnessing heterologous prime boost vaccination, with Ad5 vectored and recombinant subunit vaccines, is circumventing anti-Ad5 vector immunity induced by prime immunization [52, 53]. Due to not having to account for Ad5 vector immunity following prime immunization, along with the clinical familiarity with I.M. injection of vaccines, all mice were vaccinated intramuscularly.

**Figure 6.**
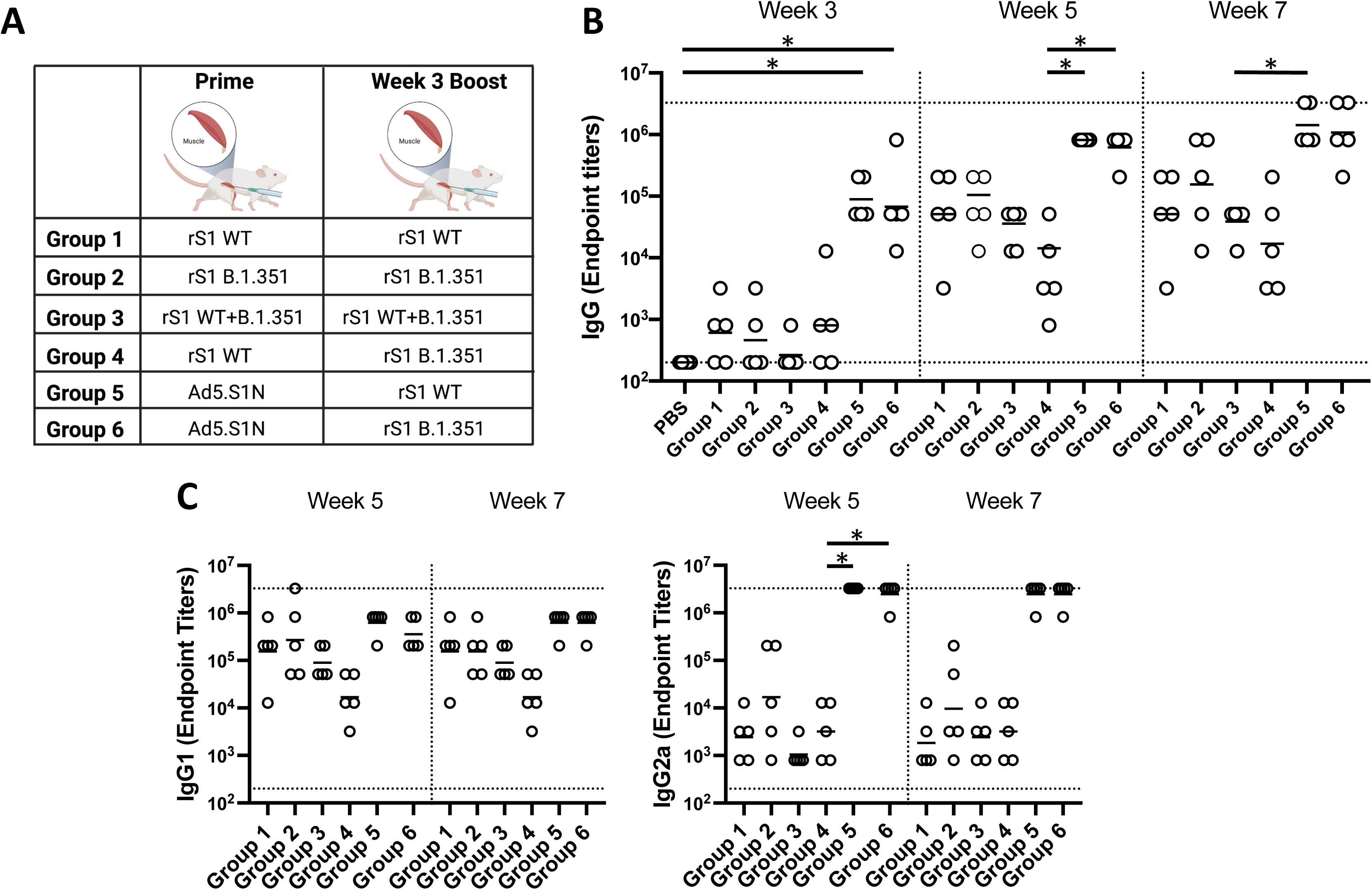
Homologous and heterologous intramuscular prime boost vaccination with variant specific recombinant S1 proteins (rS1) and Ad5.SARS-CoV-2-S1N (Ad5.SIN). BALB/c mice (n = 5 mice per groups) were immunized and boosted intramuscularly with 45ug of Wuhan rS1 (rS1 WT), South Africa rS1 (rS1 B.1.351), mixture of Wuhan rS1 and South Africa rS1 (rS1 WT+B.1351), or 1×10^10^ v.p. of Ad5.SARS-CoV-2-S1N (Ad5.S1N), while mice were immunized intramuscularly with PBS as a negative control. **(A)** Schematic layout of mice (N = 5 mice per group) vaccinations. Group 1 prime and homologous boost 15ug rS1 WT. Group 2 prime and homologous boost 15ug rS1 B.1.351. Group 3 prime and homologous boost with 15ug rS1 WT+B.1.351. Group 4 prime 15ug rS1 WT and heterologous boost 15ug rS1 B.1.351. Group 5 prime 1×10^10^ v.p. Ad5.S1N and heterologous boost 15ug rS1 WT. Group 6 prime 1×10^10^ v.p. Ad5.S1N and heterologous boost 15ug rS1 B.1.351. Antigen-specific antibody responses in mice immunized. On weeks 3,5, and 7 after vaccination, the sera from mice were collected, serially diluted (200x), and tested for the presence of SARS-CoV-2-S1-specific **(B)** IgG, **(C)** IgG1 and IgG2a antibody levels at the indicated time points by ELISA. Significance was determined by Kruskal-Wallis test followed by Dunn’s multiple comparisons (\**p* < 0.05). Horizontal lines represent geometric mean antibody titers. Horizontal dotted lines represent minimum and maximum dilutions. Results are from a single animal experiment. (N = 5 mice per group). ELISA experiments were conducted twice while neutralizing antibody experiments were conducted once.

Serum samples, collected from mice at the timepoints indicated, were used to determine S1- specific IgG, IgG1, and IgG2a endpoint titers for each immunization group using ELISA (**Fig. 6**). Heterologous Ad5.S1N prime with either WT rS1 or B.1.351 rS1 boost (Group 5 and Group 6) resulted in statistically different IgG at week 3 when compared to PBS vaccinated group (**Fig. 6B**, p < 0.05, Kruskal-Wallis test, followed by Dunn’s multiple comparisons). Groups vaccinated with rS1 protein, both WT and B.1.351 (Group 1 through Group 4), regardless of heterologous or homologous prime boost, resulted in lower IgG endpoint titers that were not statistically different from control group at week 3 and, in some cases, significantly lower than Group 5 and Group 6 at week 5 and week 7 (**Fig. 6B**, p < 0.05, Kruskal-Wallis test, followed by Dunn’s multiple comparisons). Interestingly, mice vaccinated with monomeric rS1 proteins (Group 1 through Group 4) showed a skew towards to a IgG1 dominant response rather than the balanced IgG1 and IgG2a response in Ad5.S1N primed groups (Group 5 and Group 6) (**Fig. 6C**). To evaluate the functional quality of vaccine-generated antigen-specific antibodies, we used a microneutralization assay (NT_90_) to test the ability of sera from immunized mice to neutralize the infectivity of SARS-CoV-2. The ability of sera to neutralize Beta (B.1.351) and Gamma (P.1) variants was also investigated. Sera, collected from all mice at 7 weeks after prime vaccination, were tested for the presence of SARS-CoV-2-specific neutralizing antibodies (**Fig. 7**).

**Figure 7.**
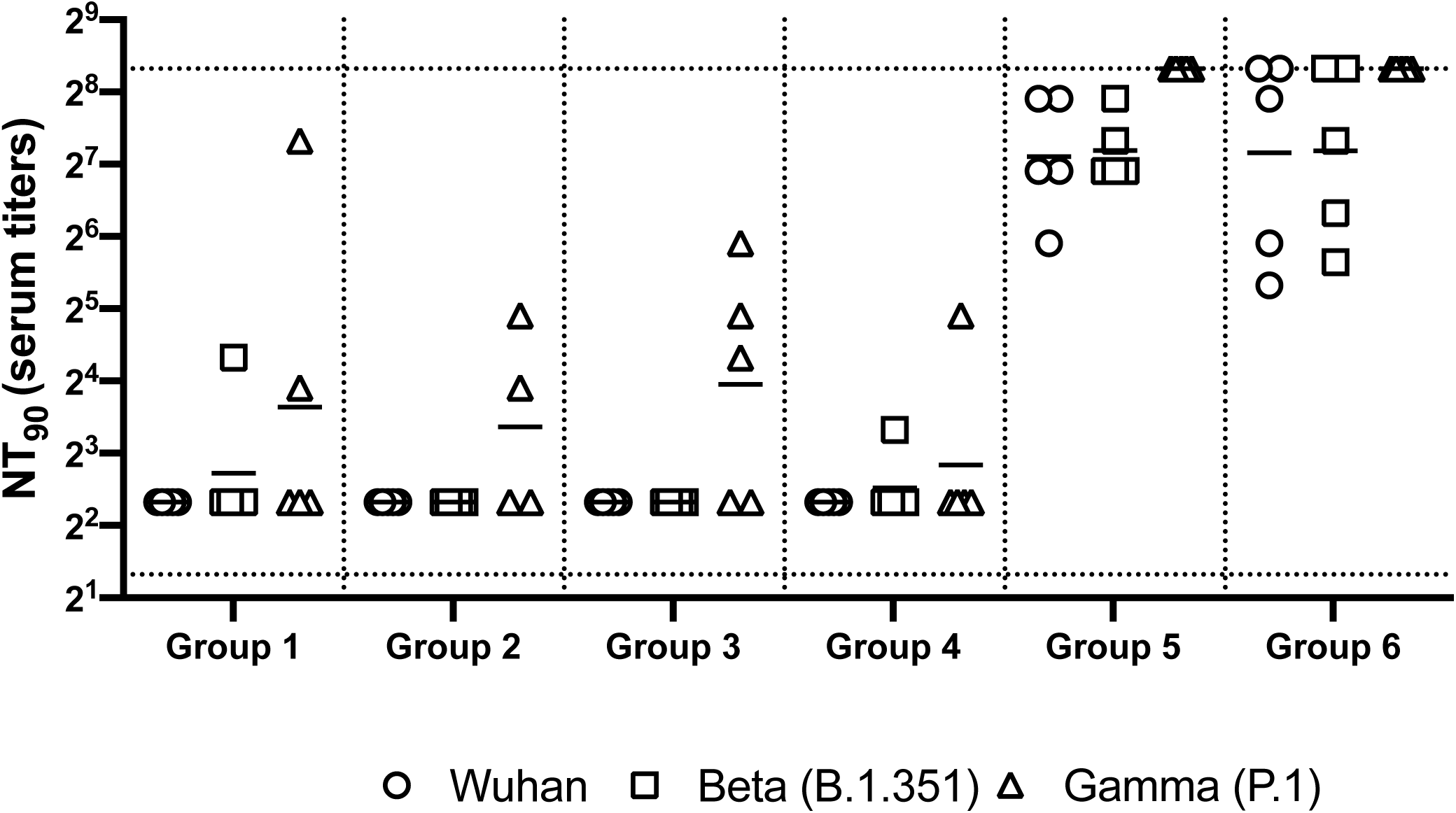
Neutralizing antibody responses in mice 7 weeks post heterologous prime-boost immunization. Group 1 prime and homologous boost 45ug rS1 WT. Group 2 prime and homologous boost 45ug rS1 B.1.351. Group 3 prime and homologous boost with 45ug rS1 WT+B.1.351. Group 4 prime 45ug rS1 WT and heterologous boost 45ug rS1 B.1.351. Group 5 prime 1×10^10^ v.p. Ad5.S1N and heterologous boost 45ug rS1 WT. Group 6 prime 1×10^10^ v.p. Ad5.S1N and heterologous boost 45ug rS1 B.1.351. Serum from immunized mice was tested for neutralizing antibodies using a plaque reduction neutralization test (PRNT) with three different SARS-CoV-2 strains from Wuhan, South Africa (Beta B.1.351), or Brazil (Gamma P.1). Neutralization of Wuhan strain represented by circle, neutralization of Beta B.1.351 represented by square, and neutralization of Gamma P.1 represented by triangle. Serum titers that resulted in a 90% reduction in SARS-CoV-2 viral plaques (NT90) compared to the virus control are reported seven weeks post initial vaccination, and bars represent geometric means (N = 5 mice per group). Results are from a single animal experiment. No neutralizing antibodies were detected in serum PBS control group (not shown). This experiment was conducted once.

Neutralizing antibodies were detected against WT, Beta, and Gamma, albeit relatively low, for monomeric rS1 vaccinated groups (Group 1 through Group 4) (**Fig. 7**). However, Ad5.S1N prime & rS1 WT boost (Group 5) and Ad5.S1N prime & rS1 B.1.351 boost (Group 6) resulted in the greatest amount of neutralizing capacity, that was not diminished against Beta (B.1.351) or Gamma (P.1). Due to the robust humoral response elicited by Group 5 and Group 6, along with the importance of long-lasting humoral response for protection against SARS-CoV-2 infection, mice were bled monthly, and on week 29 Group 1, Group 2, Group 5, and Group 6 were sacrificed to investigate the long lived S1-specfic and N-specific antibody forming cells in the bone marrow (**Fig. 8**) [57–60]. As expected, PBS vaccinated mice did not induce any S1-specfiic or N-specific antibody-producing plasma cells in the bone marrow (**Fig. 8**). Group 1, Group 2, Group 5, and Group 6 had S1-specific IgG secreting spots, with Group 5 and Group 6 having significantly higher number of spots than PBS vaccinated mice (**Fig. 8A**, p < 0.05, Kruskal-Wallis test, followed by Dunn’s multiple comparisons). Indeed, only Group 5 and Group 6 had N-specific IgG secreting spots with a significant difference from PBS, Group 1, and Group 2 mice (**Fig. 8B**, p < 0.05, Kruskal-Wallis test, followed by Dunn’s multiple comparisons).

**Figure 8.**
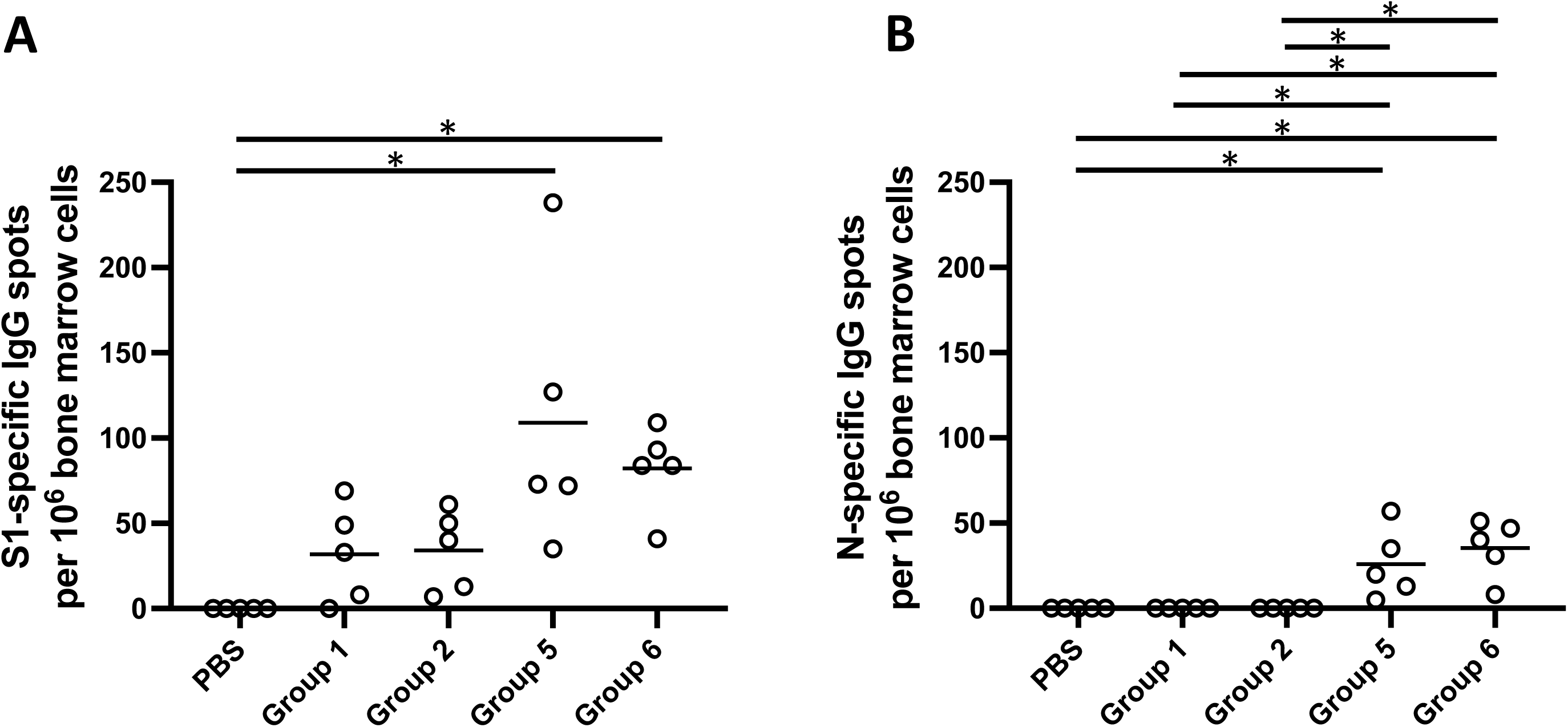
Analysis of long-term persistent antibody forming responses in the bone marrow of immunized mice. Mice were immunized as follows: Group 1 prime and homologous boost 45ug rS1 WT. Group 2 prime and homologous boost 45ug rS1 B.1.351. Group 5 prime 1×10^10^ v.p. Ad5.S1N and heterologous boost 45ug rS1 WT. Group 6 prime 1×10^10^ v.p. Ad5.S1N and heterologous boost 45ug rS1 B.1.351. Mice injected with PBS only served as negative controls. 29 weeks after immunization, S1-specific and N-specific antibody-forming cells in the bone marrow of mice were analyzed using ELISpot. **A.** Quantification of SARS-CoV-2-S1-specific antibody forming cells in the bone marrow of immunized mice. **B.** Quantification of SARS-CoV- 2 N-specific antibody forming cells in the bone marrow of immunized mice. Data are from a single experiment (n=5 per group). This experiment was conducted once.

## Discussion

More immunization programs against SARS-CoV-2 are urgently needed to battle global vaccine inequity and new viral variants. Our study presents the development, and analysis in mice, of an Ad-based COVID vaccine incorporating novel antigen design (Ad5.SARS-CoV-2- S1N). Currently approved Ad-based SARS-CoV-2 vaccines, such as Oxford-AstraZenca, Jannsen, CansinoBio, and Sputnik V COVID-19 vaccines, encode for full-length SARS-CoV-2 spike protein and are administered intramuscularly. Our vaccine encodes the gene for SARS-CoV-2-S1 subunit and SARS-CoV-2 N protein through a S1N fusion protein delivered in multiple administration routes (S.C., I.N., and I.M.). This study also investigates novel prime boost immunization strategies leveraging subunit recombinant proteins that are easy to manufacture and are relatively thermostable [61–63]. Further improvements could be achieved harnessing intracutaneous vaccination with microneedle arrays, which have been shown to deliver a wide range of recombinant DNA or protein vaccines [33, 64–66].

Our studies suggest that a single vaccination of BALB/cJ mice via either I.N. or S.C. delivery of 5×10^10^ v.p. Ad5.SARS-CoV-2-S1N was capable of inducing antigen specific IgG, Ig isotype switch, and a moderate neutralizing antibody response. S.C. delivery of Ad5.SAR-CoV-2-S1N induced a significantly increased S1-specific T cell response, when compared to S.C. delivery of AdΨ5 and S.C. delivery Ad5.SARS-CoV-2-S1. We hypothesize that this increased S1-specific T cell response to Ad5.SARS-CoV-2-S1N may be due to the inclusion of N derived T cell epitopes, in the fusion protein, aiding in processing and presentation of S1 by MHC molecules, however, this needs to be investigated further. We also illustrate that this immunogenicity can be improved by homologous prime boost strategies, using either S.C. or I.N. delivery. Particularly immunogenicity can further be improved through heterologous prime boost, with traditional I.M. injection, using subunit recombinant S1 protein. Priming with low dose (1×10^10^ v.p.) of Ad5.S1N and boosting with either WT recombinant rS1 or B.1.351 recombinant rS1 induced a robust neutralizing response, which was sustained against immune evasive variants, and a long-lived antibody forming cell response in the bone marrow 29 weeks post vaccination. Interestingly, boosting with B.1.351 recombinant rS1 did not increase the neutralizing response to SARS-CoV- 2 Beta variant virus when compared to boosting with WT recombinant rS1. The results are promising and support the use of a heterologous prime boost immunization routine using Ad5.S1N and subunit recombinant S1 protein to induce antigen specific humoral and cellular response, leading to generation of long-lived plasma cells, and a potent CD8^+^ T cell response. These results also suggest that subunit recombinant S1 protein can serve as an immunostimulatory booster and induces a robust neutralizing response to SARS-CoV-2 variants. This is critical information as COVID-19 booster demand is rapidly increasing across the globe. A booster platform using recombinant S1 protein would be thermostable, easy to manufacture, and affordable; lending it to be a preferred method to achieve global COVID-19 vaccine equity.

An important limitation regarding our findings concerning intranasal vaccination of Ad5.SARS-CoV-2-S1N is that we did not investigate key mucosal immunity aspects, such as IgA production. Our future studies will mechanistically investigate the kinetics of intranasal immunization through isolation of tissues in closer proximity to the nasal cavity. Particularly, the lack of induction of CD4^+^ and CD8^+^ T cell response to intranasal vaccination could be attributed to isolating splenocytes, rather than mucosa-associated lymph node tissue (MALT) or conducting bronchioalveolar lavage (BAL). In our future studies, we will isolate MALT and lung tissue post intranasal vaccination to investigate the localized cellular immune response post intranasal immunization. Our future studies will also include conducting BAL on intranasally immunized mice to better investigate local immunity.

We believe that the increasing CD8^+^ T cell response through including the N protein in vaccine formulation will not only help by introducing more conserved regions of SARS-CoV-2 to the immune system, potentially allowing for resistance against emerging variants, but will also assist in viral clearance. This is particularly important in the context of long covid and in populations that are at high risk of SARS-CoV-2 morbidity and mortality where the T cell response has been shown to play an important role [67–71]. While previous clinical translation of Ad vaccines has been hampered by pre-existing immunity against the Ad viral capsid, CanSino Convidicea Vaccine (Ad5-nCoV), encoding for the full S protein, was able to overcome pre-existing vector immunity while using intramuscular immunization [47, 48]. Immunization with Ad-based vaccines, such as Ad5.S1N, could be an important tool to combat COVID-19 global vaccine inequality and the emergence of immune-evasive SARS-CoV-2 variants. These finding concerning the immunogenicity of the S1N fusion antigen also adds knowledge to general COVID-19 vaccine approaches as S1N can be expressed through other vaccine vectors, such a mRNA or DNA technologies.

BALB/cJ mice have historically been used to investigate numerous different coronavirus vaccines and represent a reliable model for testing of immunogenicity of Ad5.SARS-CoV-2-S1N [33, 35, 72]. However, it is difficult to extrapolate our results in BALB/cJ mice to potential immunogenicity in humans. Particularly, our findings on the potency of S1 and N-specific T-cell response to Ad5.SARS-CoV-2-S1N vaccination may not directly translate to responses in humans due to the differences in Human Leukocyte Antigen (HLA) and mouse Major Histocompatiblity Complex (MHC). This may also explain the lack of N-specific T-cell response after vaccination, as N derived epitopes may be less able to bind to mouse MHC than S1-derived epitopes. To this effect, future studies will use more translatable animal models such as hACE2 mice and Rhesus Macaques. Our future studies will also include animal challenge models with live SARS-CoV-2 virus to investigate protection against infection and death. Along with these studies, we will also harvest axillary and cervical lymph nodes to investigate the specific kinetics and location of T-cell immunity post intramuscular vaccination.

Taken together, this study illustrates the potential of Ad5.SARS-CoV-2-S1N as it induces significant antigen specific humoral and cellular immune responses against SARS-CoV-2. Particularly, heterologous prime boost vaccine with low dose Ad5.SARS-CoV-2-S1N along with subunit recombinant S1 protein have the potential to induce a very effective virus-specific immune response against COVID-19 and emerging SARS-CoV-2 variants.

## Materials and methods

### Construction of recombinant adenoviral vectors

The coding sequence for SARS-CoV-2-S1 amino acids 1 to 661 of full-length from BetaCoV/Wuhan/IPBCAMS-WH-05/2020 (GISAID accession id. EPI_ISL_403928) flanked with SalI & BamH I-6H-Not I was codon-optimized using the UpGene algorithm for optimal expression in mammalian cells and synthesized (GenScript) ref. 2020 EBioM [73]. pAd/SARS-CoV-2-S1 was then created by subcloning the codon-optimized SARS-CoV-2-S1 gene into the shuttle vector, pAdlox (GenBank U62024), at Sal I/Not I sites. The coding sequence of N (GenBank NC_045512) having Sac I & Sal I in 5’ end and Not I & Apa I in 3’ end was synthesized and cloned in Sac I/ApaI sites in pCMV-3Tag-4A generated in pCMV3/SARS-CoV- 2-N (GenScript). For the construction of pAd/SARS-CoV-2-S1N, BamH I-6H-Not I of pAd/SARS-CoV-2-S1 was replace with Nucleoprotein gene digested with BamH I & Not I after amplified with NP-S (5’-GACGGATCCATGTCTGATAATGGACCCC-3’) & T7 (5’-TAATACGACTCACTATAGGG-3’) primers from pCMV3/SARS-CoV-2-NP. Subsequently, replication-deficient human recombinant serotype 5 adenovirus vector (Ad5.SARS-CoV-2-S1N) was generated by loxP homologous recombination and purified [35, 74, 75].

### Construction of recombinant protein expressing vectors

The coding sequence for SARS-CoV-2-S1 amino acids 1 to 661 of full-length from BetaCoV/Wuhan/IPBCAMS-WH-05/2020 (GISAID accession id. EPI_ISL_ 403928) having C-terminal tag known as ‘C-tag’, composed of the four amino acids (aa), glutamic acid – proline – glutamic acid – alanine (E-P-E-A) flanked with SalI & NotI was codon-optimized using the UpGene algorithm for optimal expression in mammalian cells and synthesized (GenScript) [73]. The construct also contained a Kozak sequence (GCC ACC) at the 5′ end. For B.1.351 variant, SARS-CoV-2-S1 mutated (Del144; K417N; E484K; N501Y; A570D; D614G) was synthesized based on above codon-optimized SARS-CoV-2-S1 from Wuhan. pAd/SARS-CoV-2-S1WU and pAd/SARS-CoV-2-SAS1SA were then created by subcloning the codon-optimized SARS-CoV-2-S1 inserts into the shuttle vector, pAdlox (GenBank U62024), at SalI/NotI sites. The plasmid constructs were confirmed by DNA sequencing.

### Transient Production of recombinant proteins in Expi293 Cells

pAd/SARS-CoV-2-S1WU and pAd/SARS-CoV-2-SAS1SA were amplified and purified using ZymoPURE II plasmid maxiprep kit (Zymo Research). For Expi293 cell transfection, we used ExpiFectamie^TM^ 293 Transfection Kit (ThermoFisher) and followed the manufacturer’s instructions. Cells were seeded 3.0 × 10^6^ cells/ml one day before transfection and grown to 4.5-5.5 × 10^6^ cells/ml. 1μg of DNA and ExpiFectamine mixtures per 1ml culture were combined and incubated for 15 min before adding into 3.0 × 10^6^ cells/ml culture. At 20h post-transfection, enhancer mixture was added, and culture was shifted to 32°C. The supernatants were harvested 5 days post transfection and clarified by centrifugation to remove cells, filtration through 0.8μm, 0.45μm, and 0.22μm filters and either subjected to further purification or stored at 4°C before purification.

### Purification of recombinant proteins

The recombinant proteins named rS1WT (Wuhan) and rS1 B.1.351 were purified using a CaptureSelect^TM^ C-tagXL Affinity Matrix prepacked column (ThermoFisher) and followed the manufacturer’s guidelines [76]. Briefly, The C-tagXL column was conditioned with 10 column volumes (CV) of equilibrate/wash buffer (20 mM Tris, pH 7.4) before sample application. Supernatant was adjusted to 20 mM Tris with 200 mM Tris (pH 7.4) before being loaded onto a 5-mL prepacked column per the manufacturer’s instructions at 5ml/min rate. The column was then washed by alternating with 10 CV of equilibrate/wash buffer, 10 CV of strong wash buffer (20 mM Tris, 1 M NaCl, 0.05% Tween-20, pH 7.4), and 5 CV of equilibrate/wash buffer. The recombinant proteins were eluted from the column by using elution buffer (20 mM Tris, 2 M MgCl_2_, pH 7.4). The eluted solution was concentrated and desalted with preservative buffer (PBS) in an Amicon Ultra centrifugal filter devices with a 50,000 molecular weight cutoff (Millipore). The concentrations of the purified recombinant proteins were determined by the Bradford assay using bovine serum albumin (BSA) as a protein standard, aliquoted, and stored at −80°C until use.

### SDS-PAGE and western blot

To evaluate the infectivity of the constructed recombinant adenoviruses, A549 (human lung adenocarcinoma epithelial cell line) cells were transduced with a multiplicity of infection of 7.5 (MOI = 7.5) of Ad5.SARS-CoV-2-S1N, Ad5.SARS-CoV-2-S1, and empty vector control (AdΨ5). At 48 hours after infection, cell supernatant was collected. The supernatants of A549 cells transduced with Ad5.SARS-CoV-2-S1N, Ad5.SARS-CoV-2-S1, and AdΨ5 were subjected to sodium dodecyl sulfate polyacrylamide gel electrophoresis (SDS-PAGE) and Western blot. Briefly, after the supernatants were boiled in Laemmli sample buffer containing 2% SDS with beta- mercaptoethanol (β-ME), the proteins were separated by Tris-Glycine SDS-PAGE gels and transferred to nitrocellulose membrane. After blocking for 1 hour at room temperature (RT) with 5% non-fat milk in TBS-T, rabbit anti-SARS-CoV spike polyclonal antibody (1:3000) (Sino Biological), or rabbit anti-SARS-CoV nucleoprotein (1:3,000) (Sino Biological) was added and incubated overnight at 4°C as primary antibody, and horseradish peroxidase (HRP)-conjugated goat anti-rabbit IgG (1:10000) (Jackson immunoresearch) was added and incubated at RT for 1 hours as secondary antibody. After washing, the signals were visualized using ECL Western blot substrate reagents and iBright 1500 (Thermo Fisher).

### Animals and immunization

For single immunization experiment, BALB/cJ mice (n = 5 animals per group in each independent experiment) were vaccinated by either I.N. delivery or S.C. injection of 5×10^10^ viral particles (v.p.) of AdΨ5 (a null Ad5 vector negative control), Ad5.SARS-CoV-2-S1, or by Ad5.SARS-CoV-2-S1N. Mice were bled from retro-orbital vein at weeks 0, 2, 4, 6, and 8 after immunization, and the obtained serum samples were diluted and used to evaluate S1-specific antibodies by enzyme-linked immunosorbent assay (ELISA). Serum samples obtained on Week 8 and 12 after vaccination were also used for microneutralization (NT) assay. For homologous and heterologous prime-boost immunization experiment, BALB/cJ mice (n = 5 animals per group in each independent experiment) were prime or boosted by either I.N. delivery or S.C. injection of 1×10^10^ viral particles (v.p.) of AdΨ5 (a null Ad5 vector negative control), Ad5.SARS-CoV-2-S1N, or by 45µg of subunit recombinant wild type S1 monomeric protein. Mice were bled from retro-orbital vein at weeks 3 and 6 after immunization, and the obtained serum samples were diluted and used to evaluate S1-specific antibodies by enzyme-linked immunosorbent assay (ELISA). Serum samples obtained on Week 6 after vaccination were also used for microneutralization (NT) assay. For heterologous prime-boost immunization experiment, BALB/cJ mice (n = 5 animals per group in each independent) were prime or boosted intramuscularly with 45µg of rS1 WT and/or rS1 B.1.351, or by Ad5.SARS-CoV-2-S1N. Mice were bled from retro-orbital vein at weeks 3, 5, and 6 after immunization, and the obtained serum samples were diluted and used to evaluate S1-specific antibodies by enzyme-linked immunosorbent assay (ELISA). Serum samples obtained on Week 6 after vaccination were also used for microneutralization (NT) assay. Mice were maintained under specific pathogen-free conditions at the University of Pittsburgh, and all experiments were conducted in accordance with animal use guidelines and protocols approved by the University of Pittsburgh’s Institutional Animal Care and Use (IACUC) Committee.

### ELISA

Sera from all mice were collected prior to immunization (week 0) and at weeks indicated after immunization and evaluated for SARS-CoV-2-S1-specific IgG, IgG1, and IgG2a antibodies using ELISA [34]. Briefly, ELISA plates were coated with 200 ng of recombinant SARS-CoV-2- S1 protein (Sino Biological) per well overnight at 4°C in carbonate coating buffer (pH 9.5) and then blocked with PBS-T and 2% bovine serum albumin (BSA) for one hour. Mouse sera were serially diluted in PBS-T with 1% BSA and incubated overnight. After the plates were washed, anti-mouse IgG-horseradish peroxidase (HRP) (1:10000, SantaCruz) was added to each well and incubated for one hour. The plates were washed three times, developed with 3,3’5,5’- tetramethylbenzidine, and the reaction was stopped. Next, absorbance was determined at 450 nm using a plate reader. For IgG1 and IgG2a ELISAs, mouse sera were diluted in PBS-T with 1% BSA and incubated overnight. After the plates were washed, biotin-conjugated IgG1 and IgG2a (1:1000, eBioscience) and biotin horseradish peroxidase (Av-HRP) (1:50000, Vector Laboratories) were added to each well and incubated for 1 hour. The plates were washed three times and developed with 3,3’5,5’-tetramethylbenzidine, the reaction was stopped, and absorbance at 450nm was determined using a plate reader.

### Flow cytometry analysis for cellular immune responses

Antigen-specific T-cell responses in the spleen of BALB/cJ mice immunized as described above were analyzed after immunization by flow cytometry, adhering to the recently published guidelines [77]. Briefly, spleens were mashed and underwent erythrocyte lysis using the Mouse Erythrocyte Lysing Kit (R&D Systems, WL2000), remaining cells were used for cellular immune response analysis. Isolated splenocytes from vaccinated and PBS control mice were stimulated with PepTivator SARS-CoV-2-S1 (a pool of S1 MHC class I– and MHC class II– restricted peptides) or SARS-CoV-2-N (a pool of N MHC class I– and MHC class II– restricted peptides) overnight in the presence of protein transport inhibitors (Golgi Stop) for the last 4 hours. Unstimulated cells were used as negative controls. Phorbol myristate acetate (PMA) and ionomycin stimulated cells served as positive controls. Cell were washed with FACS buffer (PBS, 2 % FCS), incubated with Fc Block (BD Biosciences, 553142) for 5 min at 4 °C, and stained with surface marker antibody (Ab) stain for 20 min at 4 °C. Surface Abs were used as follows: anti-CD45 (30-F11, BV480, BD Biosciences), anti-TCRb (Alexa Fluor® 700, BD Biosciences), anti-CD4 (GK1.5, BUV650, Biolegend), anti-CD8a (53-6.7, BUV570, Biolegend). For dead cell exclusion, cells were stained with Zombie NIR Fixable Viability dye (BioLegend) for 10 min at 4 °C and washed in FACS buffer. Intracellular cytokine staining (ICS) was performed on surface Ab-stained cells by first fixing and permeabilizing cells using the FoxP3 Transcription Factor Staining Buffer kit (eBioscience, 00-5523-00) following manufacturer’s instructions. Intracellular staining with anti-IFN-γ Ab (XMG1.2, BV605, Biolegend) for 30 min at 4 °C was performed. Samples were run on an Aurora (Cytek) flow cytometer and analyzed with FlowJo v10 software (BD Biosciences). Live, antigen-specific, IFN-γ producing CD8+ and CD4+ T cells were identified according to the gating strategy in Supplementary Fig. 1.

### SARS-CoV-2 microneutralization assay

Neutralizing antibody (NT-Ab) titers against SARS-CoV2 were defined according to the following protocol [78, 79]. Briefly, 50 μl of sample from each mouse, in different dilutions, were added in two wells of a flat bottom tissue culture microtiter plate (COSTAR, Corning Incorporated, NY 14831, USA), mixed with an equal volume of 100 TCID50 of a SARS-CoV2 wildtype, Beta, or Gamma variant isolated from symptomatic patients, previously titrated and incubated at 33°C in 5% CO2. All dilutions were made in EMEM (Eagle’s Minimum Essential Medium) with addition of 1% penicillin, streptomycin and glutamine and 5 γ/mL of trypsin. After 1 hour incubation at 33°C 5%CO2, 3×10^4^ VERO E6 cells [VERO C1008 (Vero 76, clone E6, Vero E6); ATCC® CRL-1586TM] were added to each well. After 72 hours of incubation at 33°C 5% CO2 wells were stained with Gram’s crystal violet solution (Merck KGaA, 64271 Damstadt, Germany) plus 5% formaldehyde 40% m/v (Carlo ErbaSpA, Arese (MI), Italy) for 30 min. Microtiter plates were then washed in running water. Wells were scored to evaluate the degree of cytopathic effect (CPE) compared to the virus control. Blue staining of wells indicated the presence of neutralizing antibodies. Neutralizing titer was the maximum dilution with the reduction of 90% of CPE. A positive titer was equal or greater than 1:5. Sera from mice before vaccine administration were always included in microneutralization (NT) assay as a negative control.

### ELISpot for antibody forming cells

The frequency of SARS-CoV-2-specific antibody producing cells in the bone marrow of mice was determined by ELISpot assay 29 weeks after immunization using our established and previously published methods [80, 81]. Briefly, 4-HBX plates were coated as described for ELISA assays, and non-specific binding was blocked with RPMI media containing 5% FCS. dilution series of cells of each individual bone marrow sample were plated in triplicates and incubated at 37 °C for 5 hours. Secondary Ab (anti-mIgG-alkaline phosphatase; Southern Biotech) was detected using 5-bromo-4-chloro-3- indolyl phosphate substrate (BCIP; Southern Biotech) in 0.5% low melting agarose (Fisher Scientific). Spots were counted using a binocular on a dissecting microscope and the detected numbers of IgG anti-SARS-CoV-2-S1 and anti-SARS-CoV-2-N antibody forming cells were calculated per million bone marrow cells.

### Statistical analysis

Statistical analyses were performed using GraphPad Prism v9 (San Diego, CA). Antibody endpoint titers and neutralization data were analyzed by Kruskal-Wallis test, followed by Dunn’s multiple comparisons, T-cell data were analyzed by one-way Welch’s ANOVA, followed by Dunnett’s T3 multiple comparisons. Significant differences are indicated by * p < 0.05. Comparisons with non-significant differences are not indicated.

## Acknowledgements

AG is funded by NIH grants UM1-AI106701, R21-AI130180, U01-CA233085) and UPMC Enterprises IPA 25565. MM is funded by NIH/NIDDK grant DK130897. These funding institutions had no role in the study design, data collection, data analysis, and interpretation of this publication. Figures 1A, 4A, and 6A were created using Biorender.com.

## Author Contributions

Conceptualization: MSK, EK, AG; Data Curation: MSK, EK, AM, FJW, EP, IC, FB, MM, AG; Formal Analysis: MSK, EK, AG, Investigation: MSK, EK, AM, FJW, SH, TWK, BLH,, EP, IC; Methodology: MSK, EK, MM, AG; Resources: MM, AG; Visualization: MSK, EK; Writing – Original Draft Preparation: MSK, EK; Writing- review and editing: MSK, EK, AM, FJW, SH, TWK, BLH, EP, IC, FB, MM, AG.

## Data availability statement

The data that support the finding of this study are available from the corresponding author upon reasonable request.

