## Supplementary Figures for "Adenovirus-Vectored SARS-CoV-2 Vaccine Expressing S1-N Fusion Protein"

### Supplementary Figure 1

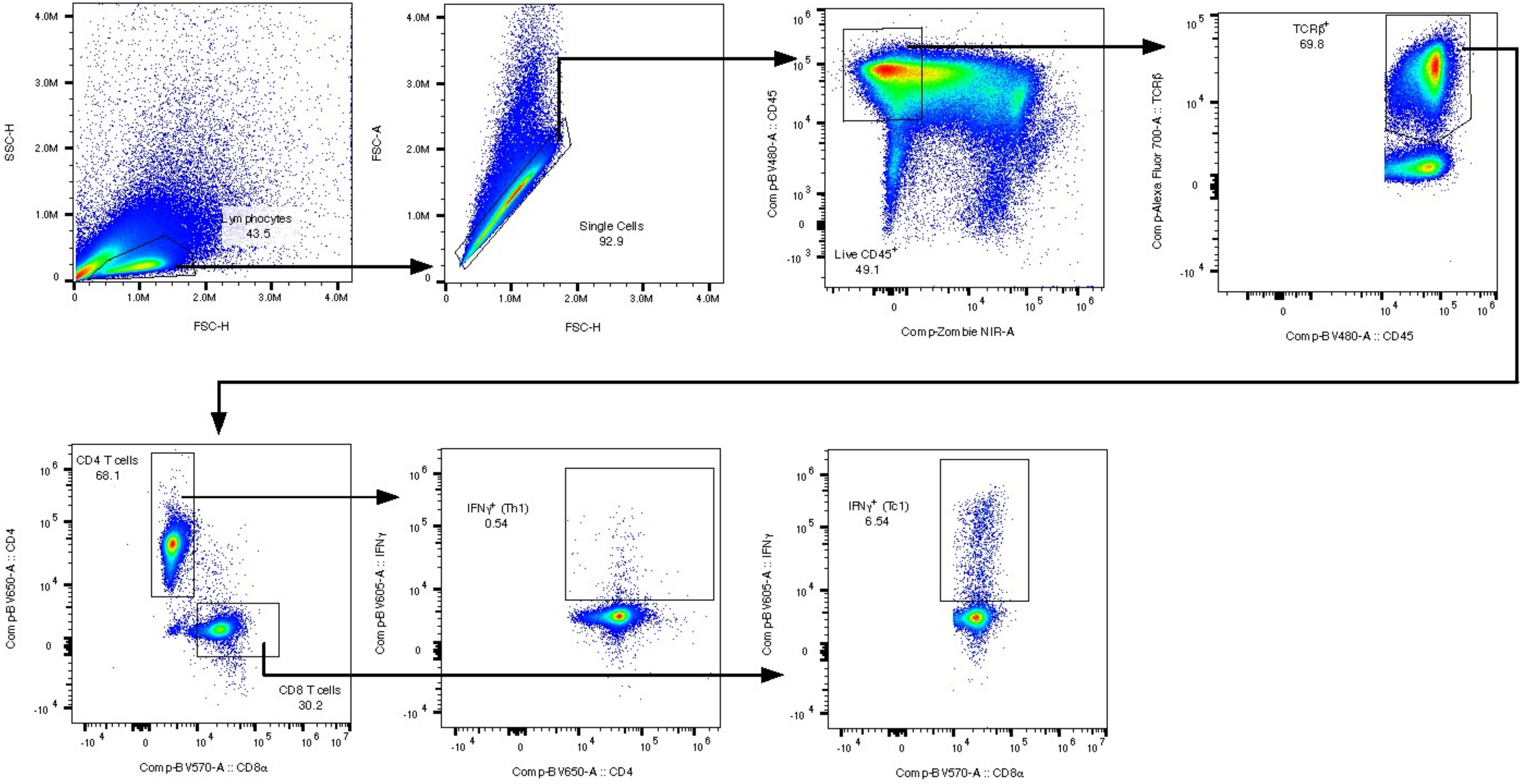


**Supplementary Figure 1. Representative gating strategy for SARS-CoV-2-S1-specific T-cell analysis.** Flow cytometry analysis was performed on splenocytes from Ad5.S1N immunized mice after ex vivo stimulation with SARS-CoV-2 S1 and intracellular cytokine staining. S1-specific live CD8+ IFN-γ+ and CD4+ IFN-γ+ were identified by singlet discrimination, exclusion of fixable viability dye (Zombie NIR–), CD45 expression, TCRβ expression, CD8a or CD4 expression, and IFN-γ expression. Black arrows indicate subsequent gating of populations, and numbers represent frequencies of cells within the indicated gate.

**Supplementary Figure 2**

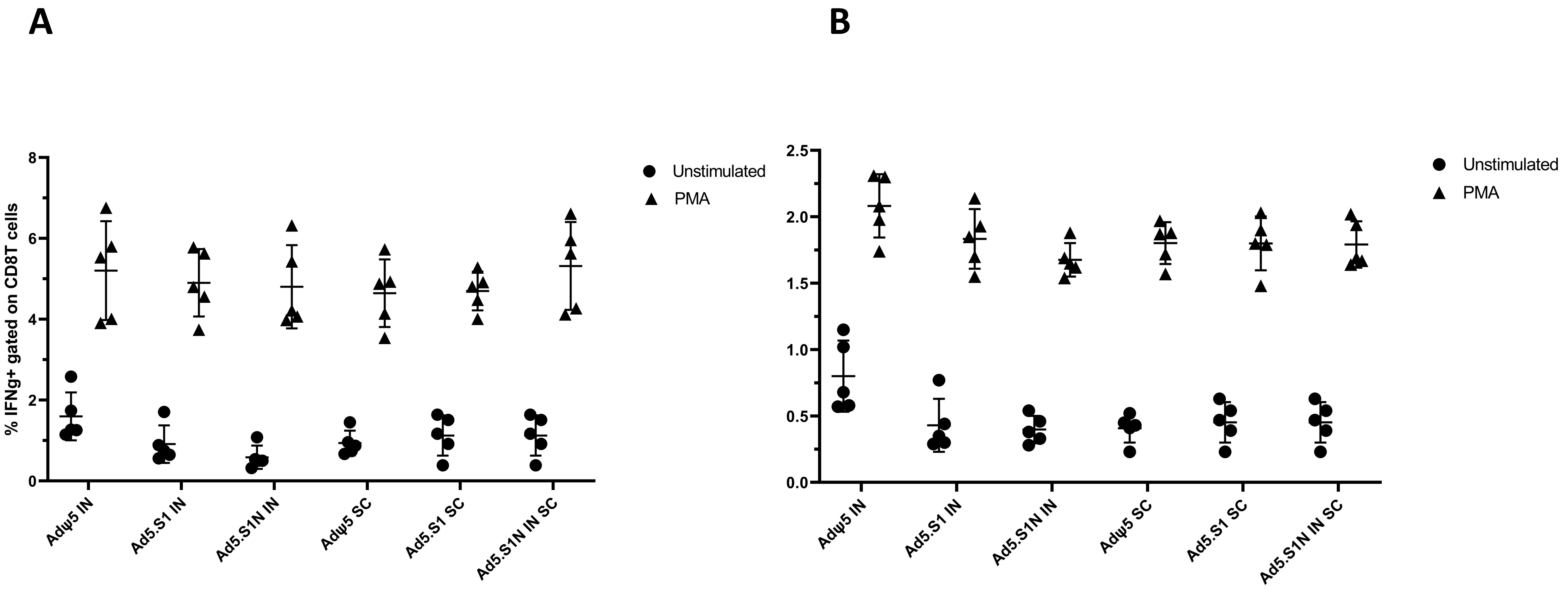


**Supplementary Figure 2. Positive and negative controls for intracellular cytokine staining (ICS) and flow cytometry. (A)** Frequencies of CD8+ IFN-$\gamma^+$ for unstimulated negative controls and PMA+Ionomycin positive controls. **(B)** Frequencies of CD4$^+$ IFN-$\gamma^+$ for unstimulated negative controls and PMA+Ionomycin positive controls. Results are mean ± SD. Unstimulated controls are represented by solid circles and PMA+Ionomycin stimulated samples are represented by solid squares. (N = 5 mice per group).

**Supplementary Figure 3**

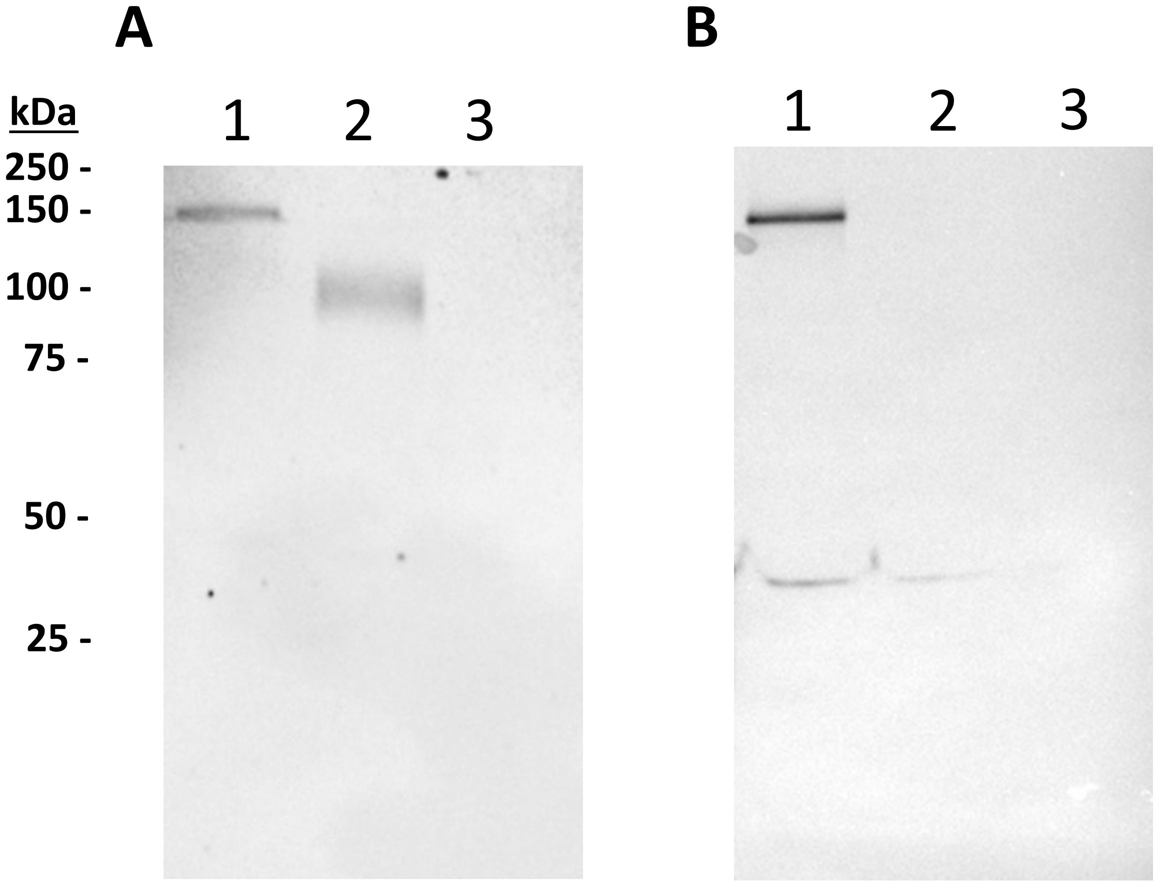


**Supplementary Figure 3. Image of original Western blots used for preparation of Fig. 1B and Fig. 1C. (A)** Detection of the SARS-CoV-2-S1N fusion protein by western blot with the supernatant of A549 cells infected with Ad5.SARS-CoV-2.S1N (Ad5.S1N) using S1 SARS-CoV-2 rabbit polyclonal antibody (lane 1). As a positive control, supernatant of A549 cells infected with Ad5.SARS-CoV-2.S1 (Ad5.S1) was loaded (lane 2). As a negative control, supernatant of A549 cells infected with an empty vector (AdΨ5) was loaded (lane 3). **(C)** Detection of the SARS-CoV-2-S1N fusion protein by western blot with the supernatant of A549 cells infected with Ad5.SARS-CoV-2.S1N (Ad5.S1N) using N SARS-CoV-2 rabbit polyclonal antibody (lane 1). As a negative control, supernatant of A549 cells infected with Ad5.SARS-CoV-2.S1 (Ad5.S1) was loaded (lane 2). As a negative control, supernatant of A549 cells infected with an empty vector (AdΨ5) was also loaded (lane 3). The supernatants were resolved on SDS-10% polyacrylamide gel after being boiled in 2% SDS sample buffer with β-ME.
